# A bifluorescent-based assay for the identification of neutralizing antibodies against SARS-CoV-2 variants of concern *in vitro* and *in vivo*

**DOI:** 10.1101/2021.06.28.450214

**Authors:** Kevin Chiem, Desarey Morales Vasquez, Jesus A. Silvas, Jun-Gyu Park, Michael S. Piepenbrink, Julien Sourimant, Michelle J. Lin, Alexander L. Greninger, Richard K. Plemper, Jordi B. Torrelles, Mark R. Walter, Juan C. de la Torre, James K. Kobie, Chengjin Ye, Luis Martinez-Sobrido

## Abstract

Severe acute respiratory syndrome coronavirus 2 (SARS-CoV-2) emerged at the end of 2019 and has been responsible for the still ongoing coronavirus disease 2019 (COVID-19) pandemic. Prophylactic vaccines have been authorized by the United States (US) Food and Drug Administration (FDA) for the prevention of COVID-19. Identification of SARS-CoV-2 neutralizing antibodies (NAbs) is important to assess vaccine protection efficacy, including their ability to protect against emerging SARS-CoV-2 variants of concern (VoC). Here we report the generation and use of a recombinant (r)SARS-CoV-2 USA/WA1/2020 (WA-1) strain expressing Venus and a rSARS-CoV-2 expressing mCherry and containing mutations K417N, E484K, and N501Y found in the receptor binding domain (RBD) of the spike (S) glycoprotein of the South African (SA) B.1.351 (beta, β) VoC, in bifluorescent-based assays to rapidly and accurately identify human monoclonal antibodies (hMAbs) able to neutralize both viral infections *in vitro* and *in vivo.* Importantly, our bifluorescent-based system accurately recapitulated findings observed using individual viruses. Moreover, fluorescent-expressing rSARS-CoV-2 and the parental wild-type (WT) rSARS-CoV-2 WA-1 had similar viral fitness *in vitro*, as well as similar virulence and pathogenicity *in vivo* in the K18 human angiotensin converting enzyme 2 (hACE2) transgenic mouse model of SARS-CoV-2 infection. We demonstrate that these new fluorescent-expressing rSARS-CoV-2 can be used *in vitro* and *in vivo* to easily identify hMAbs that simultaneously neutralize different SARS-CoV-2 strains, including VoC, for the rapid assessment of vaccine efficacy or the identification of prophylactic and/or therapeutic broadly NAbs for the treatment of SARS-CoV-2 infection.

## INTRODUCTION

The emergence of SARS-CoV-2 at the end of 2019 has been responsible for the COVID-19 pandemic ^1^. Despite numerous efforts to contain viral spread, SARS-CoV-2 disseminated worldwide and as of today it has been linked to over 175 million infections and more than 3.8 million deaths around the world ^2^. To date, one antiviral drug (remdesivir) and three human monoclonal antibodies (hMAbs) (Casirivimab/imdevimab, Bamlanivimab/etesevimab, and Sotrovimab) have been approved by the United States (US) Food and Drug Administration (FDA) for the treatment of COVID-19 ^3–5^. As of June 2021, six prophylactic vaccines against SARS-CoV-2 have been authorized by the US FDA to prevent SARS-CoV-2 infection ^6–8^. However, recent evidence suggest that newly identified SARS-CoV-2 VoC are not efficiently neutralized by sera from naturally infected or vaccinated individuals ^9^, raising concerns about the protective efficacy of current vaccines against emerging SARS-CoV-2 VoC ^10–12^.

To investigate SARS-CoV-2 infection *in vitro* and *in vivo*, including tissue and cell tropism and pathogenesis, recombinant viruses expressing a variety of reporter genes have been generated. We and others have documented the generation of recombinant (r)SARS-CoV-2 expressing fluorescent (Venus, mCherry, mNeonGreen, and GFP) or luciferase (Nluc) reporter genes ^13–16^ and their use for the identification of neutralizing antibodies (NAbs) or antivirals ^14–19^. Importantly, these reporter-expressing rSARS-CoV-2 have been shown to have growth kinetics and plaque phenotype in cultured cells similar to those of their parental rSARS-CoV-2 wild-type (WT). Current rSARS-CoV-2 have been genetically engineered to express the reporter gene replacing the open reading frame (ORF) encoding for the 7a viral protein, an approach similar to that used with SARS-CoV ^16, 20^.

Recently, we described the generation of rSARS-CoV-2 expressing reporter genes where the porcine teschovirus 1 (PTV-1) 2A autoproteolytic cleavage site was placed between the reporter gene of choice and the viral nucleocapsid (N) protein ^20^. Three major advantages of this new approach are: 1) all viral proteins are expressed (e.g. the insertion of the reporter does not replace or remove a viral protein) ^20^; 2) high levels of reporter gene expression from the N locus in the viral genome ^20^; and, 3) high genetic stability of the viral genome *in vitro* and *in vivo* because of the need of the viral N protein for genome replication and gene transcription ^20^. Importantly, this new approach allowed the visualization of infected cells *in vitro* and supported tracking SARS-CoV-2 infection *in vivo* ^20^. Notably, these new reporter-expressing rSARS-CoV-2 exhibited WT-like plaque size phenotype and viral growth kinetics *in vitro*, as well as pathogenicity in K18 human angiotensin converting enzyme 2 (hACE2) transgenic mice.

Using this strategy, we have successfully rescued Venus- and mCherry-expressing rSARS-CoV-2 USA/WA1/2020 (WA-1) and a new rSARS-CoV-2 expressing mCherry and containing mutations K417N, E484K, and N501Y present in the receptor binding domain (RBD) of the viral spike (S) glycoprotein of the South Africa (SA) B.1.351 (beta, β) VoC ^12^. Using rSARS-CoV-2 WA-1 expressing Venus and rSARS-CoV-2 SA expressing mCherry, we developed a novel bifluorescent-based assay to readily and accurately evaluate hMAbs able to specifically neutralize one or both viral variants.

Importantly, the 50% neutralizing titers (NT_50_) obtained with this new bifluorescent-based assay correlated well with those obtained using individual viruses in separated wells. Moreover, we also demonstrated the feasibility of using rSARS-CoV-2 expressing different S and fluorescent proteins (FP) to rapidly identify hMAbs able to neutralize *in vivo* both SARS-CoV-2 strains using an *in vivo* imaging system (IVIS). These new tools will help advance our understanding of efficacy of current and future SARS-CoV-2 vaccines, as well as contribute to the identification of hMAbs with broadly neutralizing activity against SARS-CoV-2 strains, including VoC, for the therapeutic or prophylactic treatment of SARS-CoV-2 infection.

## RESULTS

### Generation and characterization of rSARS-CoV-2 expressing FPs

The pBeloBAC11 plasmid encoding the full-length viral genome of SARS-CoV-2 WA-1 was used as backbone to generate the different rSARS-CoV-2 ^16, 20, 21^. We constructed new rSARS-CoV-2 reporter viruses that retained all viral genes by cloning the Venus or mCherry FP upstream of the viral N gene using the PTV-1 2A autocleavage sequence (**Figure 1A**) ^20^. Recombinant viruses expressing FPs using this experimental approach based on the use of the 2A cleavage site from the N locus do not require removing any viral genes ^20^, express higher levels of reporter gene expression compared to those previously described from the locus of the ORF7a ^20^, and are genetically more stable ^20^.

**Figure 1.**
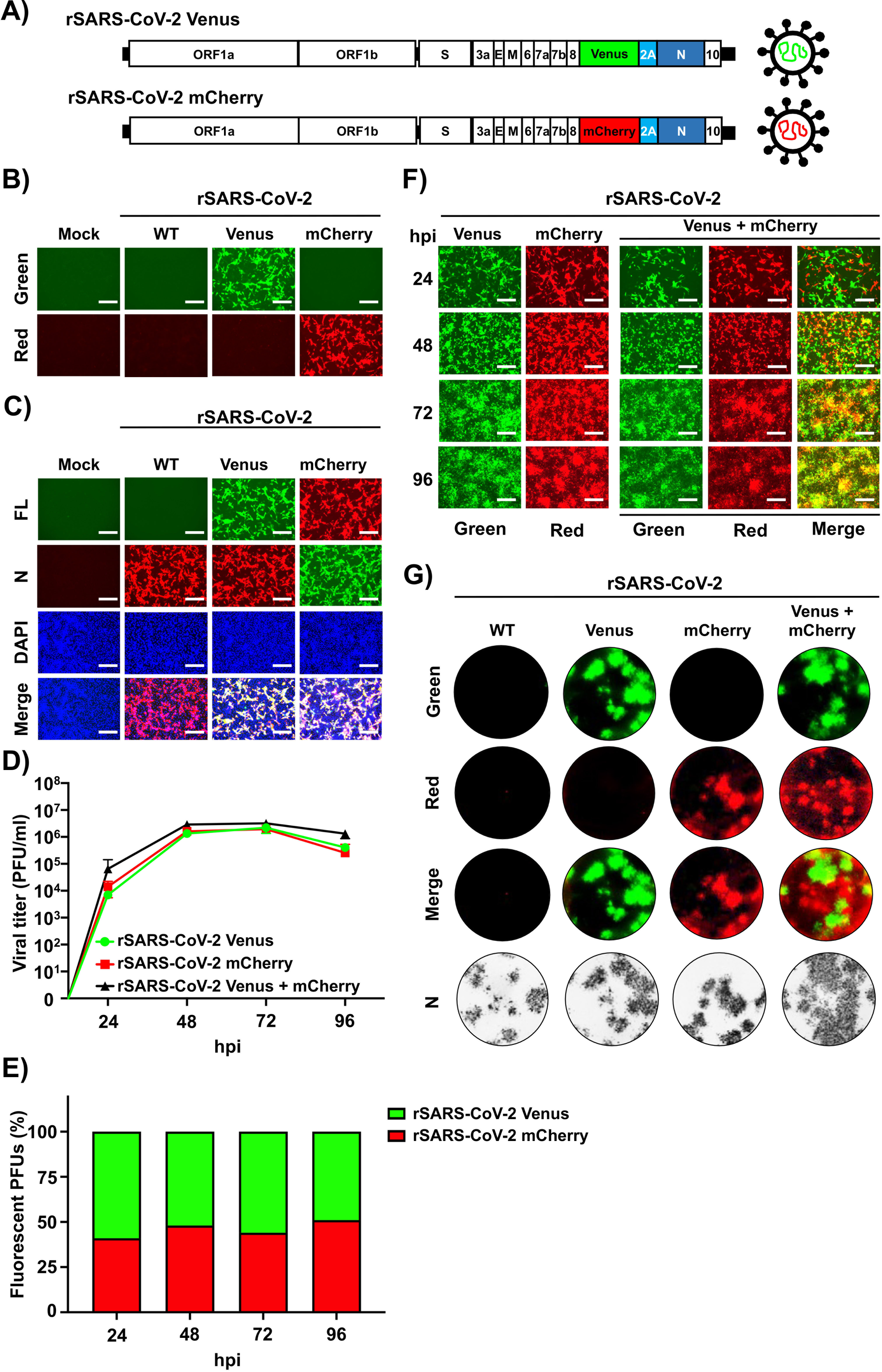
Generation and characterization of Venus and mCherry-expressing rSARS-CoV-2. A) Schematic representation of Venus and mCherry rSARS-CoV-2: Reporter genes Venus (green) or mCherry (red) were inserted upstream of the N protein (dark blue), flanked by the PTV-1 2A autocleavage sequence (light blue). B-C) Venus and mCherry expression from rSARS-CoV-2: Vero E6 cells (6-well plate format, 10^6^ cells/well, triplicates) were mock-infected or infected (MOI 0.01) with rSARS-CoV-2 Venus or rSARS-CoV-2 mCherry (**B**). At 24 hpi, cells were fixed in 10% neutral buffered formalin and visualized under a fluorescence microscope for Venus or mCherry expression. A cross-reactive mMAb against SARS-CoV N protein (1C7C7) was used for staining of infected cells (**C**). DAPI was used for nuclear staining. FL: fluorescent field. **D-F) Multi-step growth kinetics:** Vero E6 cells (6-well plate format, 10^6^ cells/well, triplicates) were mock-infected or infected (MOI 0.01) with rSARS-CoV-2 Venus and rSARS-CoV-2 mCherry, alone or together, and tissue cultured supernatants were collected at the indicated times pi to assess viral titers using standard plaque assay (**D**). The amount of Venus- and/or mCherry-positive rSARS-CoV-2 at the same times pi in cells infected with both viruses were also determined using plaque assay (**E**). Images of infected cells under a fluorescent microscope at the same times pi are shown (**F**). **G) Plaque assays:** Vero E6 cells (6-well plate format, 10^6^ cells/well, triplicates) were mock-infected or infected with ∼20 PFU of rSARS-CoV-2, rSARS-CoV-2 Venus, rSARS-CoV- 2 mCherry, or both rSARS-CoV-2 Venus and rSARS-CoV-2 mCherry. At 72 hpi, fluorescent plaques were assessed using a Chemidoc instrument. Viral plaques were also immunostained with the SARS-CoV N protein 1C7C7 cross-reactive mMAb. Fluorescent green, red and merge imaged are shown. Representative images are shown for panels B, C and F and G. Scale bars = 300 µm.

To characterize the newly generated FP-expressing rSARS-CoV-2 we first assessed Venus and mCherry expression levels. Confluent monolayers of Vero E6 cells were infected (MOI 0.01 plaque forming units (PFU)/cell) with either rSARS-CoV-2 WT, rSARS-CoV-2 Venus, rSARS-CoV-2 mCherry, or mock-infected, and then examined by fluorescence microscopy (**Figure 1B**). As expected, only cells infected with rSARS-CoV-2 Venus or rSARS-CoV-2 mCherry were detected under a fluorescent microscope (**Figure 1B**). Cells infected with rSARS-CoV-2 WT, rSARS-CoV-2 Venus, and rSARS-CoV-2 mCherry showed comparable levels of N protein expression (**Figure 1C**).

We next determined the multi-step growth kinetics of the newly generated rSARS-CoV-2. Vero E6 cells were infected (MOI 0.01 PFU/cell) with rSARS-CoV-2 Venus or rSARS-CoV-2 mCherry, individually or together, and tissue culture supernatants collected over a course of 96 h to determine viral titers (**Figure 1D**). Kinetics of production and peak titers of infectious progeny were similar for rSARS-CoV-2 expressing Venus or mCherry. Results from co-infection experiments using Venus- and mCherry-expressing rSARS-CoV-2 indicated that both viruses had similar fitness under the experimental conditions used (**Figure 1E**). This conclusion was further validated by assessing FP expression in cells infected with rSARS-CoV-2 Venus and rSARS-CoV-2 mCherry, alone or in combination (**Figure 1F**). Moreover, both rSARS-CoV-2 Venus and rSARS-CoV-2 mCherry exhibited similar plaque formation efficiency and plaque size phenotype as the parental rSARS-CoV-2 WT (**Figure 1G**).

### A bifluorescent-based assay for the identification of SARS-CoV-2 NAbs

We next assessed the feasibility of using these two FP-expressing rSARS-CoV-2, alone and in combination, to identify NAbs against SARS-CoV-2. For proof of concept, we used hMAbs 1212C2 and 1213H7, both previously shown to potently neutralize rSARS-CoV-2 ^22, 23^. The NT_50_ values of 1212C2 against rSARS-CoV-2 Venus (0.97 ng) (**Figure 2A**), rSARS-CoV-2 mCherry (1.20 ng) (**Figure 2B**), as well as rSARS-CoV-2 Venus and rSARS-CoV-2 mCherry together (0.86 ng and 0.88 ng, respectively) (**Figure 2C**) were similar to those reported using a natural SARS-CoV-2 WA-1 isolate ^16, 22^. The NT_50_ of 1213H7 against rSARS-CoV-2 Venus (2.19 ng) (**Figure 2D**), rSARS-CoV-2 mCherry (3.17 ng) (**Figure 2E**), and both, rSARS-CoV-2 Venus and rSARS-CoV-2 mCherry together (2.32 ng and 1.96 ng, respectively) (**Figure 2F**) were similar to those obtained with the natural SARS-CoV-2 WA-1 isolate ^16^. These results demonstrated the feasibility of using rSARS-CoV-2 expressing Venus and mCherry reporter genes in a new bifluorescent-based assay to identify SARS-CoV-2 NAbs.

**Figure 2.**
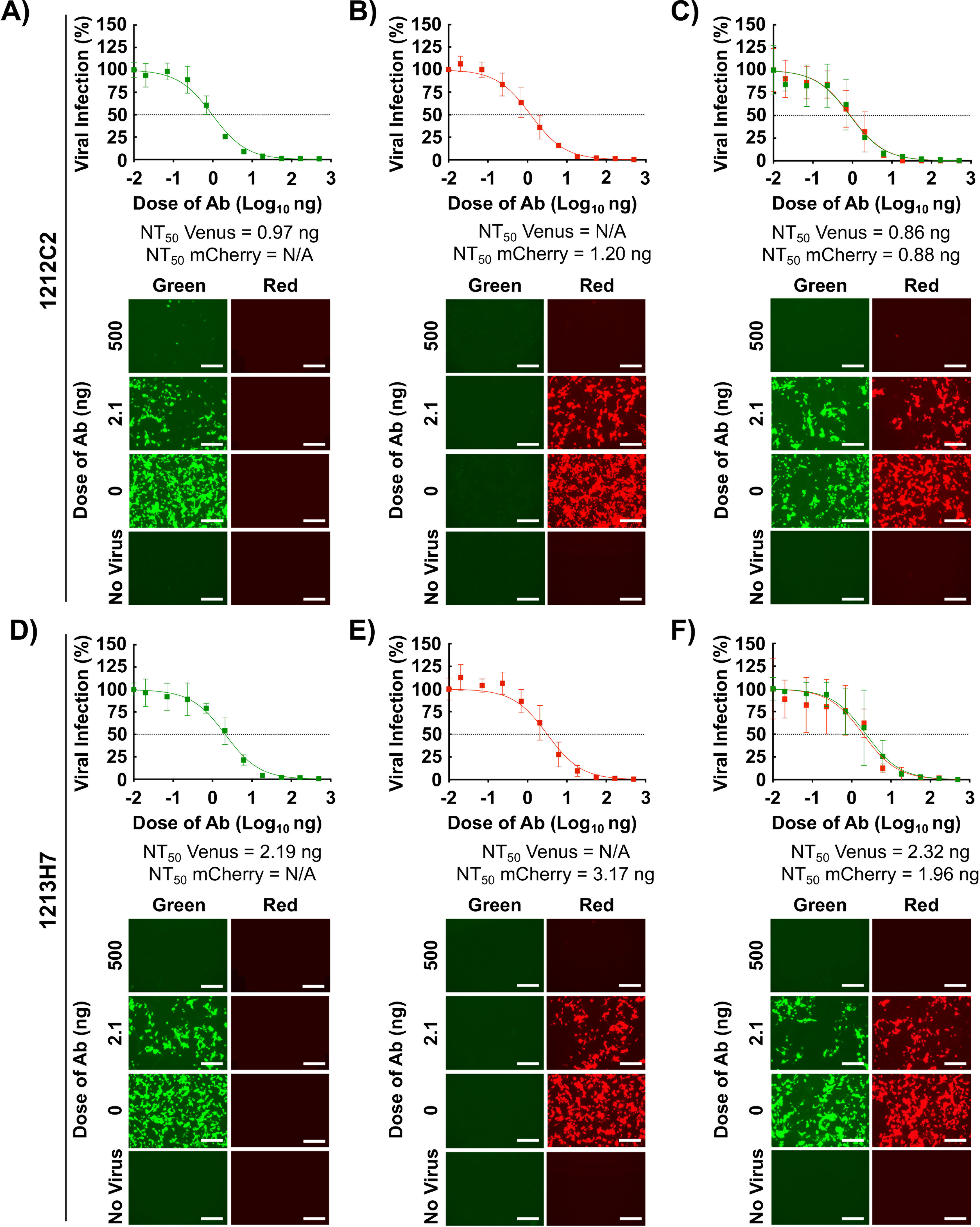
A bifluorescent-based assay to identify NAbs: Confluent monolayers of Vero E6 cells (4 x 10^4^ cells/well, 96-plate well format, quadruplicates) were infected (MOI 0.1) with rSARS-CoV-2 Venus (**A and D**), rSARS-CoV-2 mCherry (**B and E**), or both rSARS-CoV-2 Venus and rSARS-CoV-2 mCherry (**C and F**). After 1 h infection, pi media containing 3-fold serial dilutions of 1212C2 (**A-C**) or 1213H7 (**D-F**) hMAbs (starting concentration 500 ng) was added to the cells. At 48 hpi, cells were fixed with 10% neutral buffered formalin and levels of fluorescence expression were quantified in a fluorescent plate reader and analyzed using Gen5 data analysis software (BioTek). The NT_50_ values of 1212C2 and 1213H7 hMAbs for each virus, alone or in combination, were determined using GraphPad Prism. Dashed lines indicate 50% viral neutralization. Data are means and SD from quadruplicate wells. Representative images are shown. Scale bars = 300 µm.

### Generation and characterization of rSARS-CoV-2 mCherry SA

The emergence of new SARS-CoV-2 VoC, including the SA B.1.351 (beta, β) ^12^, is a major health threat since the efficacy of current vaccines against recently identified VoC may be diminished. We sought to develop an assay that would allow us to evaluate the protective efficacy of hMAbs against WA-1 and SA VoC within the same well. Towards this objective, we generated a rSARS-CoV-2 containing the K417N, E484K, and N501Y mutations found in the S RBD of the SA strain of SARS-CoV-2 and expressing also mCherry, referred to as rSARS-CoV-2 mCherry SA (**Figure 3A**). The genetic identity of the rescued rSARS-CoV-2 mCherry SA was confirmed by Sanger sequencing (**Figure 3B**).

**Figure 3.**
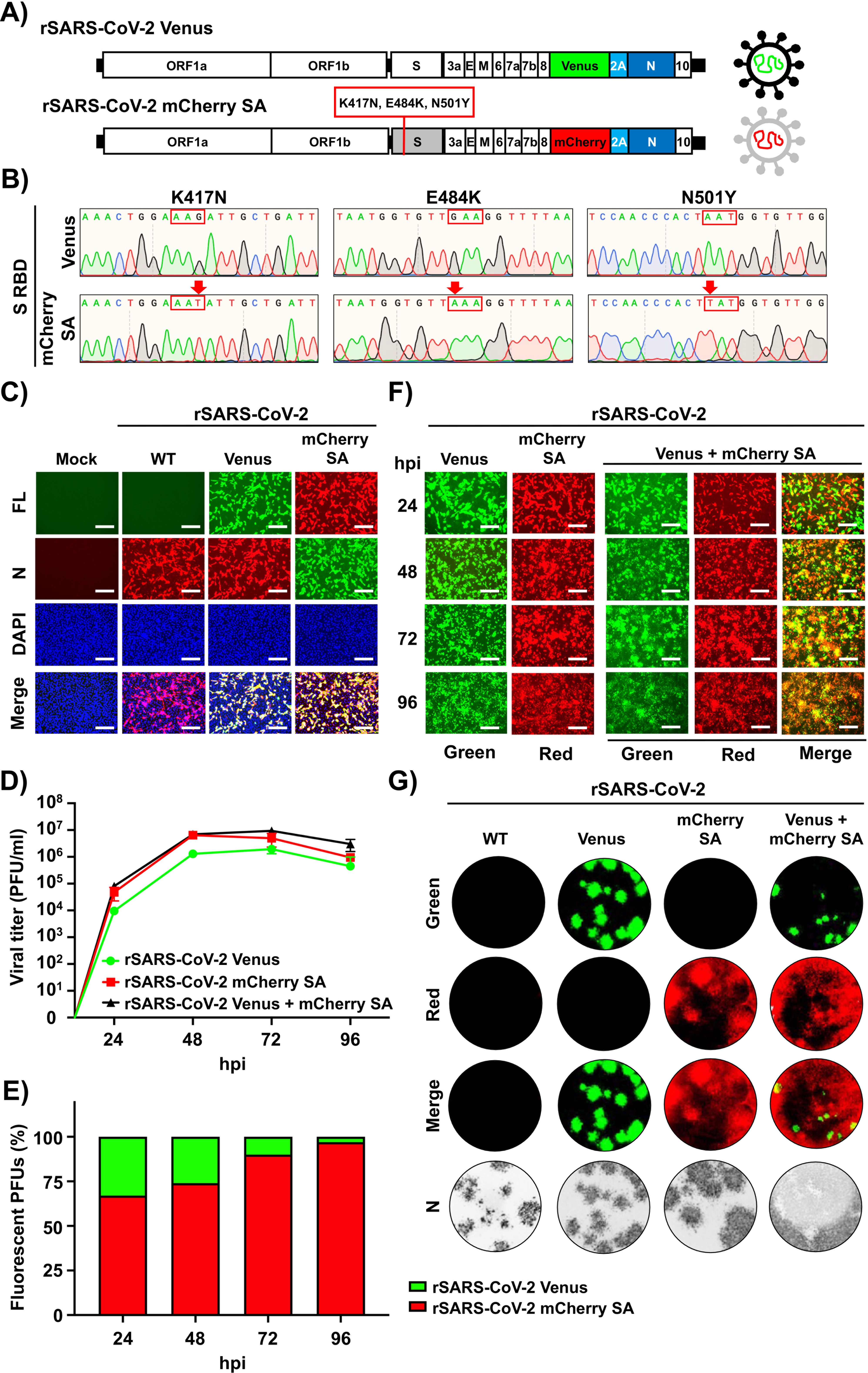
Generation and characterization of rSARS-CoV-2 mCherry SA. A) Schematic representation of rSARS-CoV-2 mCherry SA: The genome of a rSARS- CoV-2 Venus (top) and the rSARS-CoV-2 with the three mutations (K417N, E484K, and N501Y) present in the S RBD of the SA B.1.351 (beta, β) VoC expressing mCherry (bottom) is shown. **B) Sequencing of rSARS-CoV-2 mCherry SA:** Sanger sequencing results of the rSARS-CoV-2 Venus (top) and the rSARS-CoV-2 mCherry SA with the K417N, E484K, and N501Y substitutions in the RBD of the S glycoprotein (bottom) are indicated. **C) Reporter gene expression:** Vero E6 cells (6-well plate format, 10^6^ cells/well, triplicates) were mock-infected or infected (MOI 0.01) with rSARS-CoV-2, rSARS-CoV-2 Venus, or rSARS-CoV-2 mCherry SA. Infected cells were fixed in 10% neutral buffered formalin at 24 hpi and visualized under a fluorescence microscope for Venus or mCherry expression. **D-F) Multicycle growth kinetics:** Vero E6 cells (6-well plate format, 10^6^ cells/well, triplicates) were mock-infected or infected (MOI 0.01) with rSARS-CoV-2 Venus, rSARS-CoV-2 mCherry SA, or both rSARS-CoV-2 Venus and rSARS-COV-2 mCherry SA. Tissue cultured supernatants were collected at the indicated times pi to assess viral titers using standard plaque assay (**D**). The amount of Venus- and/or mCherry-positive plaques at the same times pi were determined using fluorescent microscopy (**E**). Images of infected cells under a fluorescent microscope at the same times pi are shown (**F**). **G) Plaque assays:** Vero E6 cells (6-well plate format, 10^6^ cells/well, triplicates) were mock-infected or infected with ∼20 PFU of rSARS-CoV-2, rSARS-CoV-2 Venus, rSARS-CoV-2 mCherry SA, or both rSARS-CoV-2 Venus and rSARS-CoV-2 mCherry SA. At 72 hpi, fluorescent plaques were assessed using a Chemidoc instrument. Viral plaques were also immunostained with the SARS-CoV N protein 1C7C7 cross-reactive mMAb. Fluorescent green, red and merge imaged as shown. Representative images are shown for panels C, F and G. Scale bars = 300 µm.

Next, we aimed to characterize the rSARS-CoV-2 mCherry SA by assessing reporter expression levels using rSARS-CoV-2 and rSARS-CoV-2 Venus as controls. Vero E6 cells were infected (MOI 0.01 PFU/cell) with rSARS-CoV-2 WT, rSARS-CoV-2 Venus, or rSARS-CoV-2 mCherry SA, and expression of Venus and mCherry assessed by epifluorescence microscopy (**Figure 3C**). Only cells infected with rSARS-CoV-2 Venus or rSARS-CoV-2 mCherry SA were fluorescent. However, immunostaining with the SARS-CoV cross-reactive N protein mMAb (1C7C7) detected cells infected with rSARS-CoV-2 WT, rSARS-CoV-2 Venus and rSARS-CoV-2 mCherry SA (**Figure 3C**). Next, we compared the growth kinetics of rSARS-CoV-2 mCherry SA and rSARS-CoV-2 Venus in Vero E6 cells (**Figures 3D-3F**). Interestingly, at all hpi tested, tissue culture supernatants from rSARS-CoV-2 mCherry SA infected cells had higher viral titers than those from rSARS-CoV-2 Venus infected cells (**Figure 3D**), which correlated with a higher number of mCherry than Venus positive cells in cells co-infected with rSARS-CoV-2 Venus and rSARS-CoV-2 mCherry SA **(Figures 3E and 3F**). These results were further confirmed when we assessed multiplication of rSARS-CoV-2 Venus and rSARS-CoV-2 mCherry SA by plaque assay (**Figure 3G**). Larger plaque foci were observed in cells infected with rSARS-CoV-2 mCherry SA compared to those infected with rSARS-CoV-2 Venus (**Figure 3G**). We have also observed a similar fitness advantage of a natural SARS-CoV-2 SA natural isolate over SARS-CoV-2 WA-1 ^24^.

### A bifluorescent-based assay to identify SARS-CoV-2 broadly NAbs

We next evaluated whether the rSARS-CoV-2 Venus and rSARS-CoV-2 mCherry SA could be used in a bifluorescent-based assay to identify broadly NAbs, using the 1212C2 and 1213H7 hMAbs (**Figure 2**). Preliminary data using natural SARS-CoV-2 WA-1 and SA isolates showed that 1212C2 neutralized SARS-CoV-2 WA-1 but not SARS-CoV-2 SA VoC, while 1213H7 neutralized both viral isolates ^22, 23^. As expected, 1212C2 was able to efficiently neutralize rSARS-CoV-2 Venus (NT_50_ 0.53 ng) (**Figure 4A**) but not rSARS-CoV-2 mCherry SA (NT_50_ > 500 ng) (**Figure 4B**), alone or in combination (NT_50_ 1.96 ng and > 500 ng, respectively) (**Figure 4C**). In contrast, 1213H7 was able to efficiently neutralize both rSARS-CoV-2 Venus (NT_50_ 11.89 ng) (**Figure 4D**) and rSARS-CoV-2 mCherry SA (NT_50_ 6.54 ng) (**Figure 4E**), alone or in combination (NT_50_ 12.08 and 7.97 ng, respectively) (**Figure 4F**). These results demonstrated the feasibility of using this novel bifluorescent-based assay to readily and reliably identify hMAbs with neutralizing activity against both SARS-CoV-2 strains within the same assay and that these results recapitulated those of experiments following individual viral infections and classical neutralization assays using natural viral isolates.

**Figure 4.**
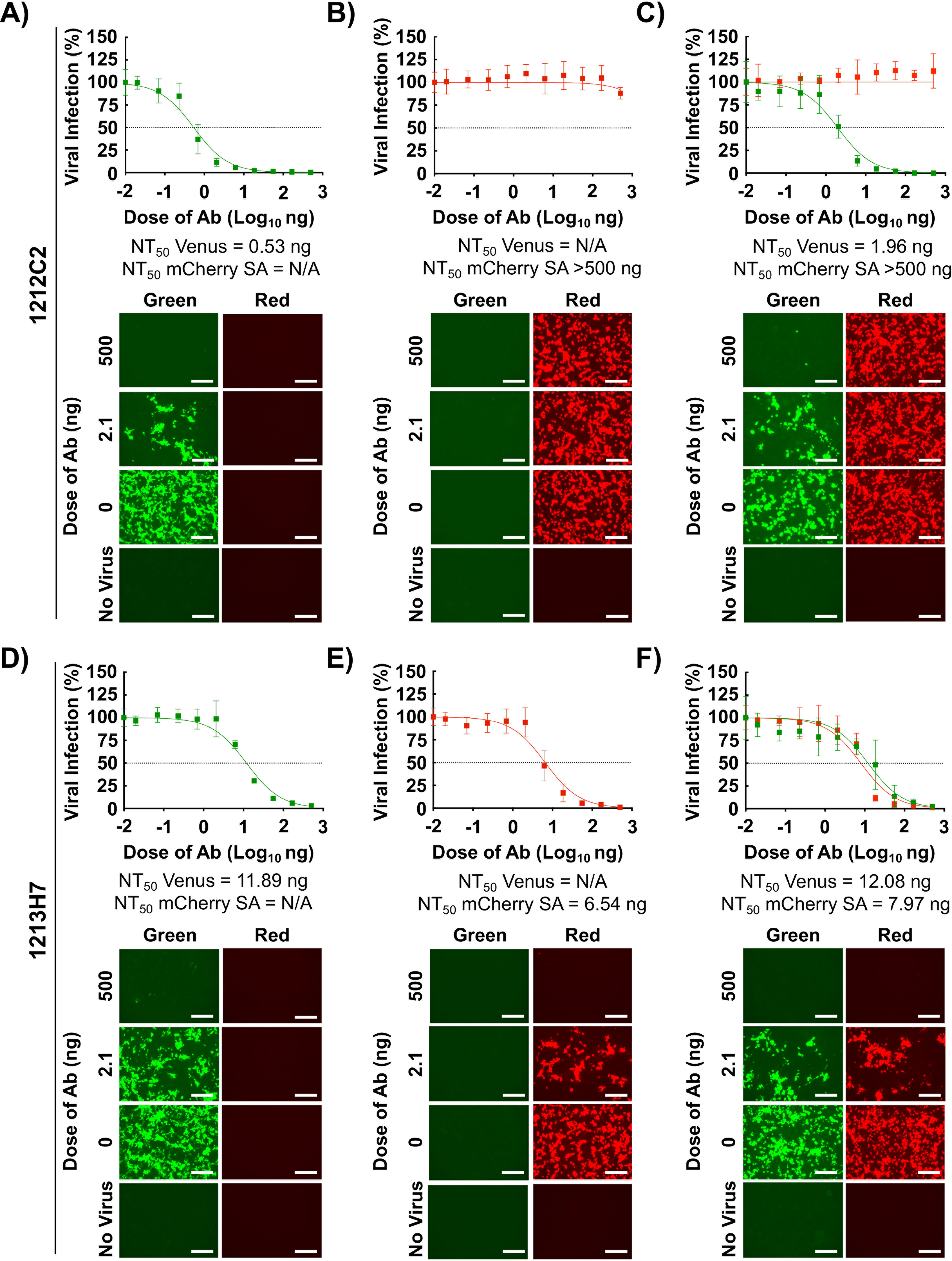
A bifluorescent-based assay to identify SARS-CoV-2 broadly NAbs: Confluent monolayers of Vero E6 cells (4 x 10^4^ cells/well, 96-plate well format, quadruplicates) were infected (MOI 0.1) with rSARS-CoV-2 Venus (**A and D**), rSARS- CoV-2 mCherry SA (MOI 0.01) (**B and E**), or both rSARS-CoV-2 Venus (MOI 0.1) and rSARS-CoV-2 mCherry SA (MOI 0.01) (**C and F**). After 1 h infection, pi media containing 3-fold serial dilutions of 12C2C2 (**A-C**) or 1213H7 (**D-F**) hMAbs (starting concentration 500 ng) was added to the cells. At 48 hpi, cells were fixed with 10% neutral buffered formalin and levels of fluorescence expression were quantified in a fluorescent plate reader and analyzed using Gen5 data analysis software (BioTek). The NT_50_ values of 1212C2 and 1213H7 hMAbs for each virus, alone or in combination, were determined using GraphPad Prism. Dashed lines indicate 50% viral neutralization. Data are means and SD from quadruplicate wells. Representative images are shown. Scale bars = 300 µm.

To further demonstrate the feasibility of this new bifluorescence-based assay to identify hMAbs able to neutralize different SARS-CoV-2 strains present in the same sample, we assessed the neutralizing activity of a selected set of previously described hMAbs ^22^. CB6, REGN10933, and REGN10987 hMAbs were used as internal controls in the assay ^25, 26^. CB6 (**Figure 5A)** and REGN10933 (**Figure 5B)** neutralized rSARS-CoV-2 Venus (NT_50_ of 1.02 and 1.53 ng, respectively) but exhibited limited (REGN10933, NT_50_ > 240.9 ng) or no (CB6, NT_50_ > 500 ng) neutralization activity against rSARS-CoV-2 mCherry SA. On the other hand, REGN10987 (**Figure 5C)** efficiently neutralized both rSARS-CoV-2 Venus and rSARS-CoV-2 mCherry SA (NT_50_ of 0.63 and 0.18 ng, respectively) (**Figure 5C**). Some of the tested hMAbs were also able to specifically neutralize rSARS-CoV-2 Venus but not rSARS-CoV-2 mCherry SA, including 1206D12 (NT_50_ 0.58 and > 500 ng, respectively) (**Figure 5D**), 1212D5 (NT_50_ 0.54 and > 500 ng, respectively) (**Figure 5E**), and 1215D1 (NT_50_ 20.31 and > 500 ng, respectively) (**Figure 5F**). We identified hMAbs with broadly neutralizing activity against both rSARS-CoV-2 Venus and rSARS-CoV-2 mCherry SA, including 1206G12 (NT_50_ of 2.23 and 1.18 ng, respectively) (**Figure 5G**), 1212F2 (NT_50_ of 31.14 and 10.64 ng, respectively) (**Figure 5H**), and 1207B4 (6.45 and 1.05 ng, respectively) (**Figure 5I**). These results support the feasibility of this novel bifluorescent-based assay to identify broad neutralizing hMAbs against different SARS-CoV-2 strains within the same assay.

**Figure 5.**
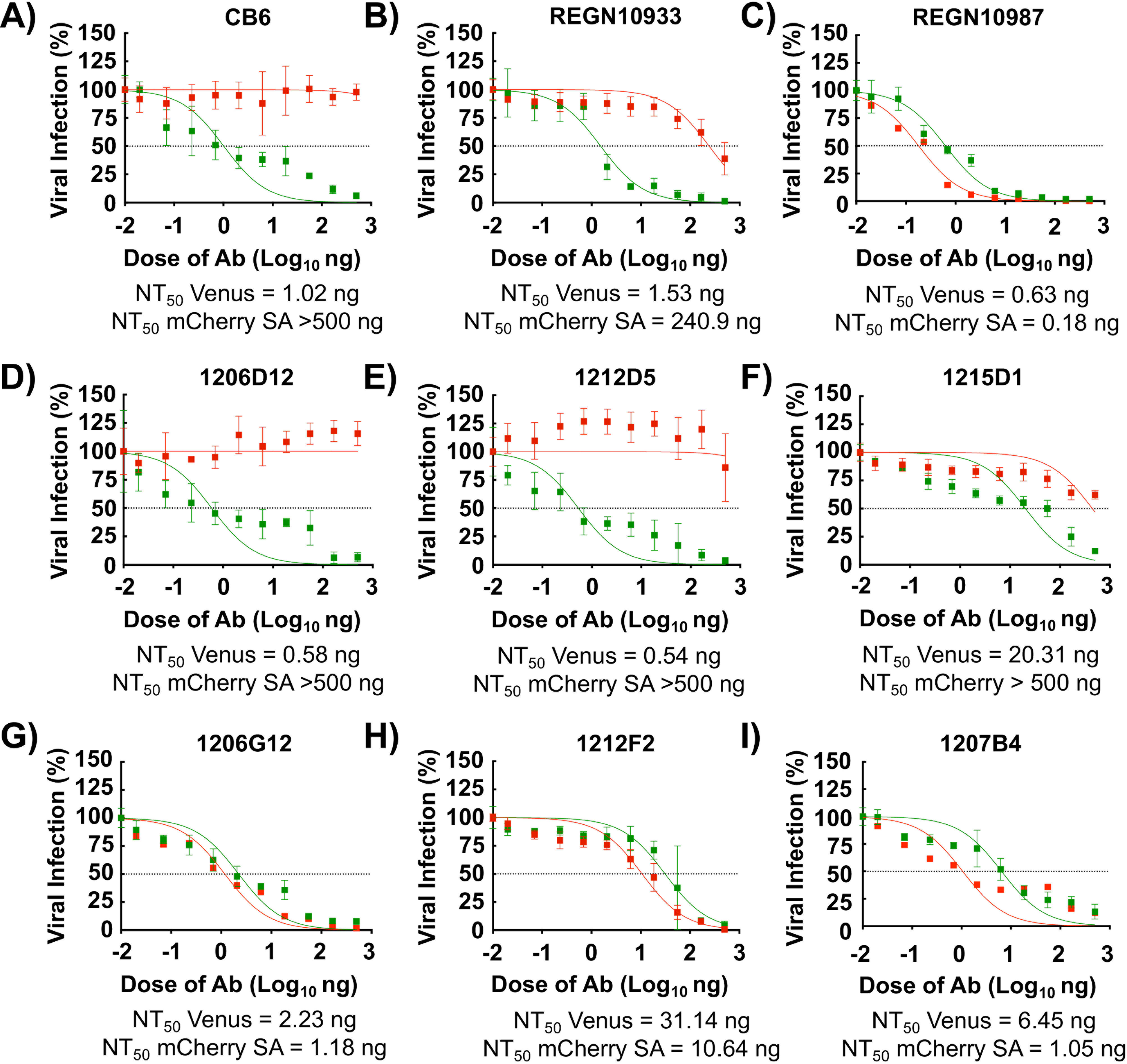
Identification of SARS-CoV-2 broadly NAbs using the bifluorescent- based assay: Confluent monolayers of Vero E6 cells (4 x 10^4^ cells/well, 96-plate well format, quadruplicates) were co-infected with rSARS-CoV-2 Venus (MOI 0.1) and rSARS-CoV-2 mCherry SA (MOI 0.01). After 1 h infection, pi media containing 3-fold serial dilutions (starting concentration 500 ng) of the indicated hMAbs was added to the cells. At 48 hpi, cells were fixed with 10% neutral buffered formalin and levels of fluorescence were quantified using a fluorescent plate reader and analyzed using Gen5 data analysis software (BioTek). The NT_50_ values of each of the hMAbs was determined using GraphPad Prism. Dashed lines indicate 50% viral neutralization. Data are means and SD from quadruplicate wells. Representative images are shown. Scale bars = 300 µm.

### An *in vivo* bifluorescent-based assay to identify SARS-CoV-2 broadly NAbs

Based on our *in vitro* results, we hypothesized that our novel bifluorescent-based assay to identify NAbs against different SARS-CoV-2 strains could be adapted to assess the neutralizing activity of hMAbs *in vivo*. To test this hypothesis, we assessed the ability of 1212C2 and 1213H7 hMAbs to neutralize rSARS-CoV-2 Venus and rSARS-CoV-2 mCherry SA, alone or in combination, in the K18 hACE2 transgenic mouse model of SARS-CoV-2 infection (**Figure 6**) ^27^. Mice were treated intraperitoneally (i.p.) with 25 mg/kg of 1212C2, 1213H7, or an IgG isotype control 24 h prior to challenge with 10^4^ PFU of rSARS-CoV-2 Venus, rSARS-CoV-2 mCherry SA, or both rSARS-CoV-2 Venus and rSARS-CoV-2 mCherry SA together. Body weight (**Figure 6A**) and survival (**Figure 6B**) were evaluated for 12 days post-infection (pi). IgG isotype control-treated mice infected with rSARS-CoV-2 Venus, rSARS-CoV-2 mCherry SA, or both rSARS-CoV-2 Venus and rSARS-CoV-2 mCherry SA together, exhibited weight loss starting on day 4 pi (**Figure 6A**) and succumbed to viral infection between days 6 to 8 pi (**Figure 6B**). However, all mice treated with 1212C2 or 1213H7 survived challenge with rSARS-CoV-2 Venus, consistent with efficient neutralization of SARS-CoV-2 WA-1 *in vitro* by these two hMAbs (**Figures 2 and 4**). In contrast, only 1213H7, but not 1212C2, was able to protect mice infected with rSARS-CoV-2 mCherry SA (**Figures 6A and 6B**), consistent with the inability of 1212C2 to neutralize rSARS-CoV-2 mCherry SA *in vitro* (**Figure 4**). When mice were co-infected with both rSARS-CoV-2 Venus and rSARS-CoV-2 mCherry SA, only mice treated with 1213H7 retained their initial body weight and survived infection (**Figures 6A and 6B**, right panels), similar to results obtained using individual infections.

**Figure 6.**
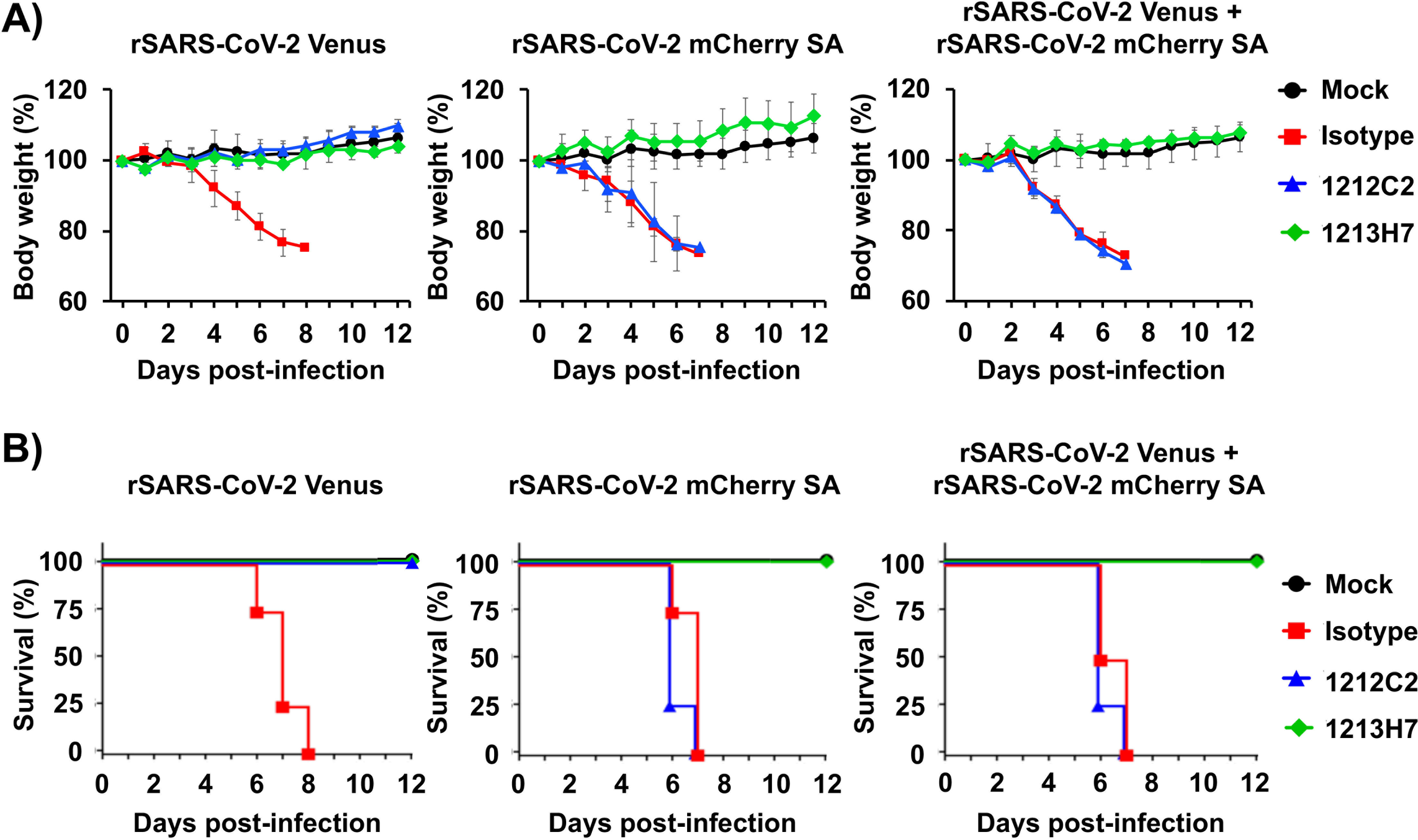
Prophylactic activity of 1212C2 and 1213H7 against rSARS-CoV-2 Venus and rSARS-CoV-2 mCherry SA, alone or in combination, in K18 hACE2 transgenic mice: Six-to-eight-week-old female K18 hACE2 transgenic mice (n=5) were treated (i.p.) with 25 mg/kg of IgG isotype control, hMAb 1212C2, or hMAb 1213H7 and infected with 10^4^ PFU of rSARS-CoV-2 Venus (left), rSARS-CoV-2 mCherry SA (middle) or both, rSARS-CoV-2 Venus and rSARS-CoV-2 mCherry SA (right). Mice were monitored for 12 days for changes in body weight (**A**) and survival (**B**). Data represent the means and SD of the results determined for individual mice.

### Use of FP expression to assess kinetics of SARS-CoV-2 multiplication in the lungs of infected K18 hACE2 transgenic mice

We next examined whether FP expression could be used as a surrogate of SARS-CoV-2 multiplication in the lungs of infected mice, providing a readout to assess the *in vivo* protective activity of 1212C2 and 1213H7 hMAbs through IVIS (**Figure 7**). K18 hACE2 transgenic mice were treated (i.p., 25 mg/kg) with IgG isotype control, 1212C2, or 1213H7 hMAbs, 24 h before infection (10^4^ PFU/mouse) with rSARS-CoV-2 Venus and/or rSARS-CoV-2 mCherry SA, singly or in combination. Mock-infected mice were included as control. At days 2 and 4 pi, Venus and mCherry expression in the lungs was evaluated using IVIS (**Figure 7A**) and quantified using Aura imaging software (**Figure 7B**). Excised lungs were also evaluated in a blinded manner by a certified pathologist to provide gross pathological scoring (**Figure 7A**). Both Venus and mCherry expression were detected in the lungs of mice treated with the IgG isotype control and infected with rSARS-CoV-2 Venus and/or rSARS-CoV-2 mCherry SA, respectively (**Figure 7A**), alone or in combination. Fluorescent signal increased from day 2 to day 4 pi in the lungs of all IgG isotype control-treated infected mice (**Figure 7B**). Mice treated with 1212C2 and infected with rSARS-CoV-2 Venus showed no detectable Venus signal, indicating that 1212C2 protects against rSARS-CoV-2 Venus infection (**Figure 7A**, top panel). In contrast, 1212C2-treated mice infected with rSARS-CoV-2 mCherry SA expressed mCherry in the lungs (**Figure 7A**, middle panel). In mice treated with 1212C2 and co-infected with both rSARS-CoV-2 Venus and rSARS-CoV-2 mCherry SA, we observed only mCherry expression, consistent with the ability of 1212C2 to neutralize rSARS-CoV-2 Venus but not rSARS-CoV-2 mCherry SA (**Figure 7A**, bottom panel).

**Figure 7.**
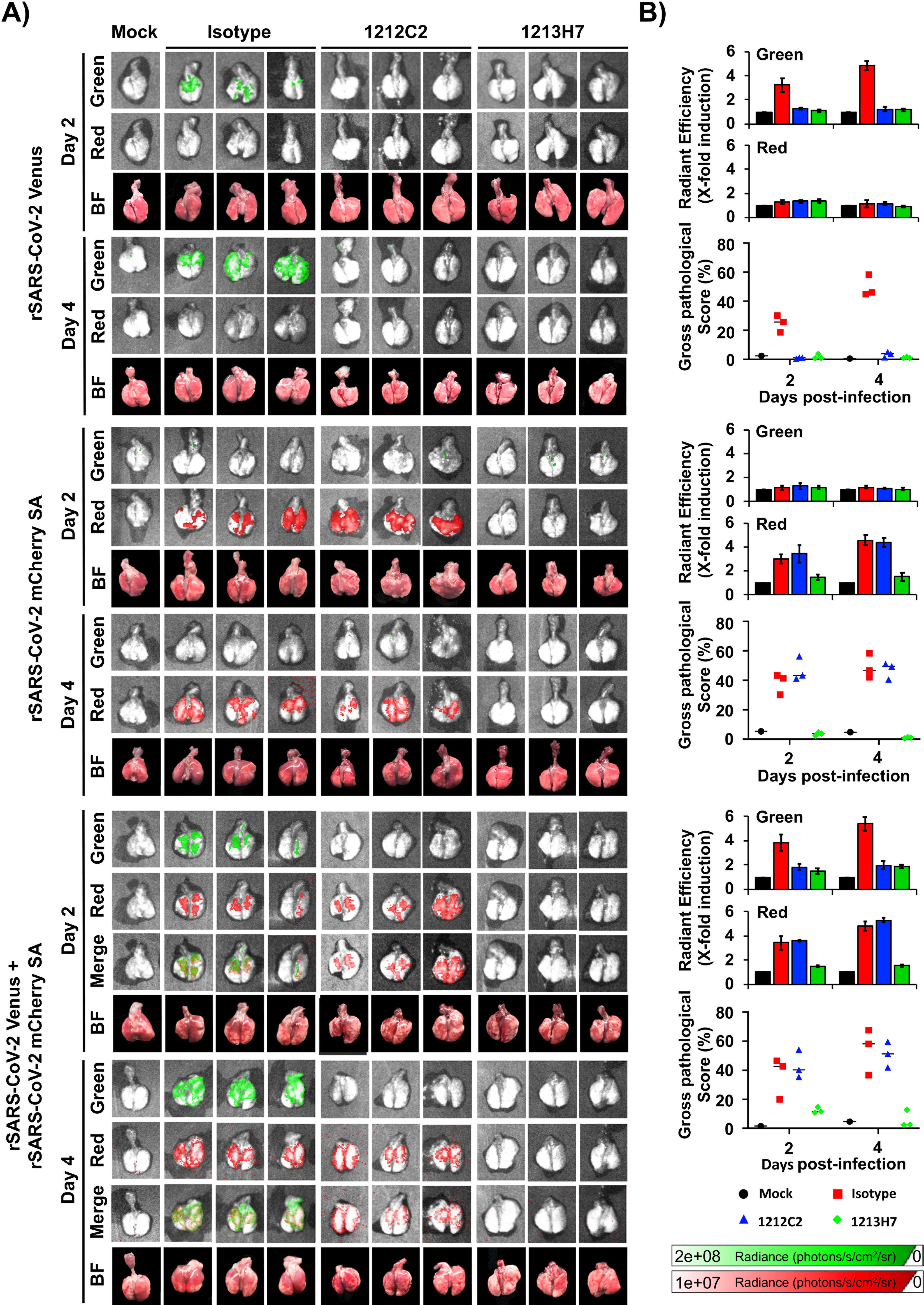
Kinetics of fluorescent expression in the lungs of K18 hACE2 transgenic mice treated with 1212C2 or 1213H7 hMAbs and infected with rSARS-CoV-2 Venus and rSARS-CoV-2 mCherry SA: Six-to-eight-week-old female K18 hACE2 transgenic mice (n=3) were injected (i.p.) with 25 mg/kg of an IgG isotype control, hMAb 1212C2, or hMAb 1213H7 and infected with 10^4^ PFU of rSARS-CoV-2 Venus (top), rSARS-CoV-2 mCherry SA (middle), or both, rSARS-CoV-2 Venus and rSARS-CoV-2 mCherry SA (bottom). At days 2 and 4 pi, lungs were collected to determine Venus and mCherry fluorescence expression using an Ami HT imaging system (**A**). BF, bright field. Venus and mCherry radiance values were quantified based on the mean values for the regions of interest in mouse lungs (**B**). Mean values were normalized to the autofluorescence in mock-infected mice at each time point and were used to calculate fold induction. Gross pathological scores in the lungs of mock-infected and rSARS-CoV- 2-infected K18 hACE2 transgenic mice were calculated based on the % area of the lungs affected by infection.

Corroborating our previous *in vitro* and *in vivo* results (**Figures 4 and 6**, respectively), mice treated with 1213H7 were protected against infection with both rSARS-CoV-2 Venus and rSARS-CoV-2 mCherry SA, when administered alone or in combination, and presented no detectable fluorescence in the lungs (**Figure 7A**). These data were further supported by quantification of the average radiant efficiency of fluorescence signals, which were high in the lungs of IgG isotype control-treated mice infected with rSARS-CoV-2 Venus or rSARS-CoV-2 mCherry SA, and in the lungs of 1212C2-treated mice infected with rSARS-CoV-2 mCherry SA (**Figure 7B**). Importantly, gross pathological scoring correlated with levels of FP expression in the lungs of infected mice.

As predicted, IgG isotype control-treated K18 hACE2 transgenic mice infected with rSARS-CoV-2 Venus (**Figure 8A**), rSARS-CoV-2 mCherry SA (**Figure 8B**), or both rSARS-CoV-2 Venus and rSARS-CoV-2 mCherry SA (**Figure 8C**) presented high viral titers. In contrast, lungs of 1212C2-treated and infected mice had undetectable levels of rSARS-CoV-2 Venus (**Figure 8A**), but high titers of rSARS-CoV-2 mCherry SA when mice were individually infected (**Figure 8B**) or co-infected with both viruses (**Figure 8C**). In 1213H7-treated and infected mice, we did not detect rSARS-CoV-2 Venus (**Figure 8A**) or rSARS-CoV-2 mCherry SA (**Figure 8B**), including double infected mice (**Figure 8C**), consistent with the ability of 1213H7 to potently neutralize both viruses *in vitro* and *in vivo* (**Figures 4 and 6**, respectively), Lung homogenates from IgG isotype control-treated mice infected with both reporter viruses contained ∼25% and ∼75% of Venus and mCherry, respectively, positive plaques by day 2 pi. This finding suggested that rSARS-CoV-2 mCherry SA had a higher fitness than rSARS-CoV-2 Venus *in vivo* (**Figure 8D**), which was similar to our *in vitro* studies (**Figure 3**). Notably, by day 4 pi all viral plaques were mCherry-positive, further supporting a higher fitness of rSARS-CoV-2 mCherry SA compared to rSARS-CoV-2 Venus *in vivo* (**Figure 8D**). Lung homogenates from 1212C2-treated mice contained rSARS-CoV-2 mCherry SA, reflecting the ability of 1212C2 to efficiently neutralize rSARS-CoV-2 Venus but not rSARS-CoV-2 mCherry SA. In contrast, no viral plaques were detected in lung homogenates from mice treated with 1213H7, as this hMAb efficiently neutralizes both viruses. We obtained similar results in the nasal turbinate (**Figure 8**, middle) and brain (**Figure 8**, bottom) of hMAb-treated and infected K18 hACE2 transgenic mice.

**Figure 8.**
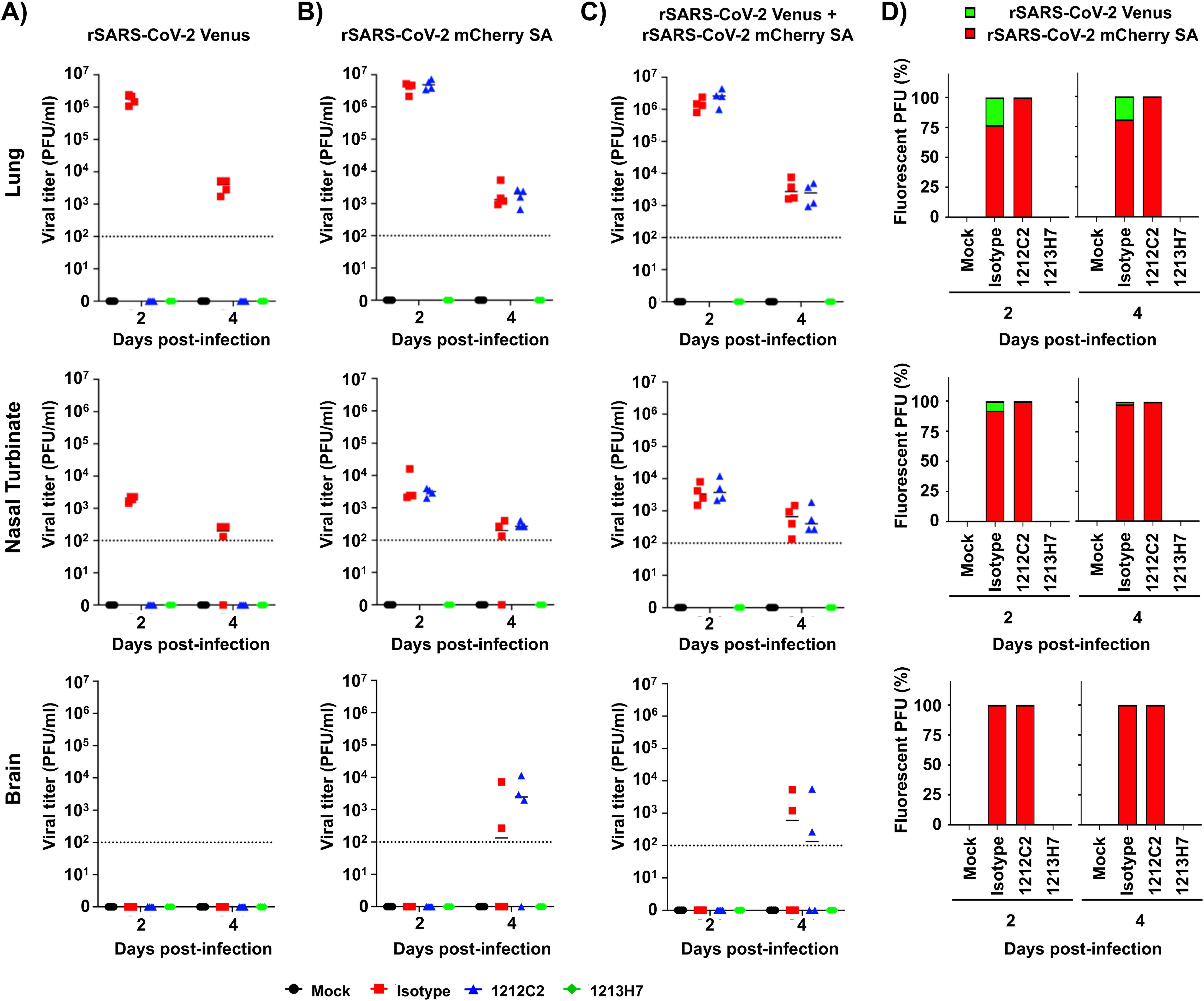
Viral titers in the lungs, nasal turbinate and brain of K18 hACE2 transgenic mice treated with 1212C2 or 1213H7 hMAbs and infected with rSARS- CoV-2 Venus and rSARS-CoV-2 mCherry SA: Six-to-eight-week-old female K18 hACE2 transgenic mice (n=3) injected (i.p.) with 25 mg/kg of an IgG isotype control, hMAb 1212C2, or hMAb 1213H7, and infected with 10^4^ PFU of rSARS-CoV-2 Venus (**A**), rSARS-CoV-2 mCherry SA (**B**), or both rSARS-CoV-2 Venus and rSARS-CoV-2 mCherry SA (**C**). Viral titers in the lungs (top), nasal turbinate (middle) and brain (bottom) at days 2 and 4 pi were determined by plaque assay in Vero E6 cells. Bars indicates the mean and SD of lung virus titers. Dotted lines indicate the limit of detection. **D)** Quantification of rSARS-CoV-2 Venus and rSARS-CoV-2 mCherry SA in the lungs (top), nasal turbinate (middle) and brain (bottom) from mice co-infected with both rSARS-CoV-2 Venus and rSARS-CoV-2 mCherry SA at days 2 and 4 pi.

## DISCUSSION

The COVID-19 pandemic caused by SARS-CoV-2 started at the end of 2019 ^1^.

Despite US FDA-authorized prophylactic vaccines ^6–8^ and some, although still limited, available therapeutic antiviral drugs (remdesivir) and neutralizing hMAbs (Casirivimab/imdevimab, Bamlanivimab/etesevimab, and Sotrovimab) interventions ^3–5^, over 175 million infections and more than 3.8 million deaths have been attributed to the COVID-19 pandemic globally ^2^. As with other viruses, SARS-CoV-2 is continuously evolving, resulting in viral variants (e.g. VoC) that are becoming dominant within the human population due to increased fitness, transmission and/or resilience against naturally or vaccine induced immune responses. To date, several VoC have been identified, including the United Kingdom B.1.1.7 (alpha, α) ^28^, SA B.1.351 (β) ^12^, Brazil P.1 (gamma, γ) ^29, 30^, India B.1.617.2 (delta, δ) ^31^, and California B.1.427 (epsilon, ε) ^32^. There is limited information on the ability of current vaccines to protect against these newly identified SARS-CoV-2 VoC ^9, 10, 33^. Moreover, it is likely that additional VoC will emerge in the future.

Reporter-expressing recombinant viruses can circumvent limitations imposed by the need for secondary methods to detect the presence of viruses in infected cells. These reporter viruses have been used to evaluate viral infections, identify therapeutics, and to study viral virulence *in vivo*. Here, we have documented the generation of novel rSARS-CoV-2 to facilitate tracking infection of two different SARS-CoV-2 strains (WA-1 and SA) *in vitro* and *in vivo* based on the use of two different FPs (Venus and mCherry, respectively). The FP-expressing rSARS-CoV-2 encode the fluorescent Venus or mCherry proteins from the locus of the N protein, without the need for deletion of any viral protein ^20^. Notably, the use of this approach to generate FP-expressing rSARS-CoV-2 resulted in higher FP expression levels than those afforded by rSARS-CoV-2 expressing FPs from the locus of the viral ORF7a protein ^20^. Moreover, rSARS-CoV-2 expressing reporter genes from the N locus are more genetically stable than those expressing reporter genes from the ORF7a locus of the SARS-CoV-2 genome ^20^.

We showed that rSARS-CoV-2 expressing Venus or mCherry from the N locus exhibited similar growth kinetics, peak titers, and plaque phenotype as the parental WT rSARS-CoV-2 WA-1 strain. Importantly, we were able to use these novel reporter rSARS-CoV-2 in bifluorescent-based assays to determine the neutralization efficacy of hMAbs based on FP expression levels. We also generated rSARS-CoV-2 mCherry SA, an mCherry-expressing rSARS-CoV-2 containing the K417N, E484K, and N501Y mutations in the RBD of the S glycoprotein of the SA VoC. Notably, rSARS-CoV-2 mCherry SA had higher fitness than rSARS-CoV-2 Venus in cultured cells, as evidenced by higher viral titers reached and a bigger plaque size phenotype.

Interestingly, when used in the bifluorescent-based assay, hMAb 1212C2 was unable to neutralize rSARS-CoV-2 mCherry SA, but efficiently neutralized rSARS-CoV-2 Venus. In contrast, hMAb 1213H7 displayed efficient neutralization of both rSARS-CoV-2 Venus and rSARS-CoV-2 mCherry SA. Importantly, these *in vitro* results correlated with *in vivo* studies in which K18 hACE2 transgenic mice pre-treated with 1212C2 were protected against challenge with rSARS-CoV-2 Venus but not rSARS-CoV-2 mCherry SA, alone or in combination. In contrast, mice treated with 1213H7 were protected against lethal challenge with both reporter-expressing rSARS-CoV-2, alone or in combination. These protection results were corroborated through IVIS studies, in which fluorescence and viral titers demonstrated the neutralizing protective efficacy of 1212C2 against rSARS-CoV-2 Venus but not rSARS-CoV-2 mCherry SA, while 1213H7 efficiently protected mice against challenge with both viruses, alone or in combination. These results prove the feasibility of using both rSARS-CoV-2 Venus and rSARS-CoV-2 mCherry SA to accurately assess the ability of hMAbs to efficiently neutralize one or both SARS-CoV-2 strains, alone or in combination, *in vitro* and/or *in vivo*, and establish that the readouts of the bifluorescent-based assays correlate well with those of individual viral infections.

rSARS-CoV-2 expressing FP or luciferase reporter genes have been described by us and others ^13–16^, but in this study we have documented, for the first time, the use of two rSARS-CoV-2 expressing different FP and S glycoproteins in a bifluorescent-based assay to identify NAbs exhibiting differences in their neutralizing activity against different SARS-CoV-2 strains present in the same biological sample *in vitro* and *in vivo*. Notably, this approach can be extended to identify broadly NAbs against different SARS-CoV-2 VoC by generating rSARS-CoV-2 expressing additional FP and containing the S glycoproteins of different VoC in multiplex-based fluorescent assays *in vitro* and/or *in vivo*. These reporter rSARS-CoV-2 expressing the S glycoprotein of VoC also represent an excellent option to investigate viral infection, dissemination, pathogenesis and therapeutic interventions, including protective efficacy of vaccines or antivirals, for the treatment of SARS-CoV-2 infection in cultured cells and/or in validated animals models of SARS-CoV-2 infection.

## METHODS

### Biosafety

Experiments involving the use of infectious SARS-CoV-2 were performed at biosafety level 3 (BSL3) containment laboratories at Texas Biomedical Research Institute. All experiments using SARS-CoV-2 were approved by the Institutional Biosafety Committee (IBC) at Texas Biomedical Research Institute.

### Cells

African green monkey kidney epithelial cells (Vero E6, CRL-1586) were grown and maintained in Dulbecco’s modified Eagle’s medium (DMEM) supplemented with 10% fetal bovine serum (FBS) and 1X PSG (100 units/ml penicillin, 100 µg/ml streptomycin, and 2 mM L-glutamine), and incubated at 37°C in an 5% CO_2_ atmosphere.

### Generation of pBeloBAC11-SARS-CoV-2 encoding fluorescent proteins (FP)

The pBeloBAC11 plasmid (NEB) containing the entire viral genome of SARS-CoV-2 USA/WA1/2020 (WA-1) isolate (accession no. MN985325) has been described ^20, 21^.

The rSARS-CoV-2 expressing Venus or mCherry from the locus of the viral N protein using the PTV-1 2A autocleavage sequence were generated as previously described ^20, 21^. The rSARS-CoV-2 containing mutations K417N, E484K, and N501Y present in the receptor binding domain (RBD) within the spike (S) gene of the South African (SA) B.1.351 (beta, β) VoC ^12^ and expressing mCherry was generated using standard molecular biology techniques. Plasmids containing the full-length genome of the different rSARS-CoV-2 were analyzed by digestion using specific restriction enzymes and validated by deep sequencing. Oligonucleotides for cloning the Venus or mCherry FP, or K417N, E484K, and N501Y mutations, are available upon request.

### Generation of rSARS-CoV-2 expressing FP

Wild-type (WT, WA-1), Venus (Venus WA-1), and mCherry (mCherry WA-1) reporter-expressing rSARS-CoV-2, as well as rSARS-CoV-2 encoding the SA B.1.351 (beta, β) mutations K417N, E484K, and N501Y in the S RBD expressing mCherry (mCherry SA) were rescued as previously described ^21, 34^. Briefly, confluent monolayers of Vero E6 cells (1.2 x 10^6^ cells/well, 6-well plate format, triplicates) were transfected with 4 µg/well of pBeloBAC11-SARS-CoV-2 (WA-1), -2A/Venus, -2A/mCherry, or - 2A/mCherry-SA-RBD plasmids using Lipofectamine 2000 (Thermo Fisher). After 24 h post-transfection, media was exchanged with post-infection (pi) media (DMEM containing 2% FBS), and 24 h later cells were scaled up to T75 flasks and incubated for 72 h at 37°C. Viral rescues were first confirmed under a brightfield microscope by assessing cytopathic effect (CPE) before supernatants were collected, aliquoted, and stored at -80°C. To confirm the rescue of rSARS-CoV-2, Vero E6 cells (1.2 x 10^6^ cells/well, 6-well plates, triplicates) were infected with virus-containing tissue culture supernatants and incubated at 37°C in a 5% CO_2_ incubator for 48 h. Viruses were detected by fluorescence or immunostaining with a SARS-CoV N protein cross reactive mouse (m)MAb (1C7C7). Plaque assays were used to determine viral titers (plaque forming units, PFU)/ml). Viral stocks were generated by infecting fresh monolayers of Vero E6 cells at low multiplicity of infection (MOI, 0.0001) for 72 h before aliquoted and stored at -80°C.

### Sequencing

To confirm the identity of the rescued rSARS-CoV-2 mCherry SA, total RNA from infected (MOI 0.01) Vero E6 cells (1.2 x 10^6^ cells/well, 6-well format, triplicates) was extracted using TRIzol reagent (Thermo Fisher Scientific), and used in RT-PCR reactions to amplify a fragment of 1,174 bp around the RBD of the S gene. RT-PCR was done using SuperScript II reverse transcriptase (Thermo Fisher Scientific) and Expand high-fidelity PCR system (Sigma-Aldrich). RT-PCR products were purified on 0.7% agarose gel and subjected to Sanger sequencing (ACGT). All primer sequences are available upon request.

### Immunofluorescence assays

Confluent monolayers of Vero E6 cells (1.2 x 10^6^ cells/well, 6-well format, triplicates) were mock-infected or infected (MOI 0.01) with WT, Venus-, or mCherry-expressing rSARS-CoV-2 WA-1 (WA-1, Venus WA-1, or mCherry WA-1, respectively); or rSARS-CoV-2 mCherry SA. At 48 hours post-infection (hpi), cells were submerged in 10% neutral buffered formalin at 4°C overnight for fixation and viral inactivation, and then permeabilized with 0.5% Triton X-100 phosphate-buffered saline (PBS) at room temperature for 10 min. Thereafter, cells were washed with PBS before blocking with 2.5% bovine albumin serum (BSA) PBS for 1 h. Cells were then incubated with 1 µg/ml of SARS-CoV anti-N mMAb 1C7C7 in 1% BSA at 37°C for 1 h. Reporter-expressing rSARS-CoV-2 were detected directly by epifluorescence and using either Alexa Fluor 594 goat anti-mouse IgG (Invitrogen; 1:1,000) or fluorescein isothiocynate (FITC)- conjugated goat anti-mouse IgG (Dako; 1:200), depending on whether the viruses express Venus or mCherry, respectively. Cell nuclei were detected with 4’, 6’-diamidino- 2-phenylindole (DAPI, Research Organics). An EVOS M5000 imaging system was used to acquire representative images (10X magnification).

### Viral growth kinetics

Vero E6 cells (1.2 x 10^6^ cells/well, 6-well plate format, triplicates) were infected (MOI 0.01) at 37°C for 1 h. After viral adsorption, cells were washed with PBS and incubated at 37°C in pi media. At 24, 48, 72, and 96 hpi, fluorescence-positive cells were imaged with an EVOS M5000 fluorescence microscope for rSARS-CoV-2 expressing Venus or mCherry FP, and viral titers in the tissue culture supernatants were determined by plaque assay and immunostaining using the anti-SARS-CoV N mMAb 1C7C7. Mean values and standard deviation (SD) were calculated with Microsoft Excel software.

### Plaque assays and immunostaining

Confluent monolayers of Vero E6 cells (2 x 10^5^ cells/well, 24-well plate format, triplicates) were infected with WT or reporter-expressing rSARS-CoV-2 for 1 h before being overlaid with pi media containing 1% agar (Oxoid) and incubated at 37°C in a 5% CO_2_ incubator. After 72 h, cells were fixed in 10% neutral buffered formalin overnight at 4°C. Next, overlays were removed, PBS was added to each well, and fluorescent plaques were detected and quantified using a ChemiDoc MP imaging system (Bio-Rad). Cells were then permeabilized with 0.5% Triton X-100 in PBS for 5 min, blocked with 2.5% BSA in PBS for 1 h, and incubated with the SARS-CoV N mMAb 1C7C7, and plaques detected using a Vectastain ABC kit and DAB HRP Substrate kit (Vector laboratories) following the manufacturers’ instructions.

### A bifluorescence-based neutralization assay

The hMAbs used in this study were generated and purified as described ^22^. CB6, REGN10987 and REGN10933 hMAbs were included as controls ^25, 26^. To test the neutralizing activity of hMAbs, confluent monolayers of Vero E6 cells (4 x 10^4^ cells/well, 96-plate well format, quadruplicates) were infected (MOI of 0.01 or 0.1) with the indicated rSARS-CoV-2 for 1 h at 37°C. After viral absorption, pi media containing 3-fold dilutions of the indicated hMAbs (starting concentration of 500 ng/well) were added to the cells and incubated at 37°C for 48 h. Cells were then fixed in 10% neutral buffered formalin overnight and washed with PBS, before fluorescence signal was measured and quantified using a Synergy LX microplate reader and Gen5 data analysis software (Bio- Tek). The mean and SD of viral infections were calculated from individual wells of three independent experiments conducted in quadruplicates with Microsoft Excel software.

Non-linear regression curves and NT_50_ values were determined using GraphPad Prism Software (San Diego, CA, USA, Version 8.2.1). Representative images were captured with an EVOS M5000 Imaging system (Thermofisher) at 10X magnification.

### Mouse experiments

All animal protocols were approved by Texas Biomed IACUC (1718MU). Six-to-eight-week-old female K18 human angiotensin converting enzyme 2 (hACE2) transgenic mice were purchased from The Jackson Laboratory and maintained in the Animal Biosafety Laboratory level 3 (ABSL-3) at Texas Biomedical Research Institute. All mouse procedures were approved by Texas Biomedical Research Institute IACUC. To assess the *in vivo* efficacy of hMAbs, K18 hACE2 transgenic mice (n=5/group) were anesthetized with isoflurane and injected (i.p.) with hMAbs IgG isotype control, 1212C2 or 1213H7 (25 mg/kg) using a 1 ml syringe 23-25 gauge 5/8 inch needle 24 h prior to challenge with rSARS-CoV-2. For viral challenges, mice were anesthetized and inoculated intranasally (i.n.) with 10^4^ plaque forming units (PFU) of the indicated rSARS-CoV-2 and monitored daily for morbidity as determined by changes in body weight, and survival. Mice that lost greater than 25% of their initial weight were considered to have reached their experimental endpoint and were humanely euthanized. In parallel, K18 hACE2 transgenic mice (n=3/group) were treated (i.p.) with 1212C2 or 1213H7 hMAbs and challenged i.n. with 10^4^ PFU of the indicated rSARS-CoV-2 for viral titer determination. Viral titers in the lungs of infected mice at days 2 and 4 pi were determined by plaque assay. *In vivo* fluorescence imaging of mouse lungs was conducted using an Ami HT *in vivo* imaging system, IVIS (Spectral Instruments). Mice were euthanized with a lethal dose of Fatal-Plus solution and lungs were surgically extracted and washed in PBS before imaging in the Ami HT. Images were analyzed with Aura software to determine radiance with the region of interest (ROI), and fluorescence signal was normalized to background signal of lungs from mock-infected mice. Bright field images of lungs were captured using an iPhone X camera. After imaging, lungs were homogenized using a Precellys tissue homogenizer (Bertin Instruments) in 1 ml of PBS and centrifuged at 21,500 x g for 10 min to pellet cell debris. Clarified supernatants were collected and used to determine viral titers by plaque assay. Macroscopic pathological scoring was determined from the percent of total surface area affected by congestion, consolidation, and atelectasis of excised lungs, using NIH ImageJ software as previously described ^21, 35^.

### Statistical analysis

All data are presented as mean values and SD for each group and were analyzed using Microsoft Excel software. A two-tailed Student *t* test was used to compare the means between two groups. A *P* value of less than 0.05 (*P*<0.05) was considered statistically significant.

### Data availability

All of the data supporting the findings of this work can be found within the paper.

The raw data are available from the corresponding authors upon request.

## ACKNOWLEDGEMENTS

We are grateful to Dr. Thomas Moran at The Icahn School of Medicine at Mount Sinai for providing the SARS-CoV cross-reactive 1C7C7 N protein mMAb.

## AUTHOR CONTRIBUTIONS

C.Y. rescued the rSARS-CoV-2 expressing FP; K.C. conducted the *in vitro* experiments; D.M.V. conducted the bifluorescent neutralization assays; K.C. J.P., J.S., and D.M.V. conducted the *in vivo* experiments; J. J. K., M.S.P., and M. R. W. provided critical reagents; J.C.T., J.S., M.J.L., A.L.G., and R.K.P. deep sequenced the viruses; K.C. and D.M.V. drafted the manuscript; J.B.T. provide support for the *in vitro* and *in vivo* studies at the BSL3 and ABSL3 facilitates, respectively, at Texas Biomedical Research Institute. L.M-S. conceived the study, revised, and finalized the manuscript. All authors review and approve the manuscript.

## COMPETING INTERESTS

J.-G.P., M.S.P., M.R.W., J.J.K., and L.M.-S. are co-inventors on a patent that includes claims related to some of the hMAbs described in the manuscript.

